# Identification of salt tolerant genotypes in wheat using stress tolerance indices

**DOI:** 10.1101/2021.02.15.431202

**Authors:** Behnam Bakhshi, Seyed Mohammad Taghi Tabatabaei, Mohammad Reza Naroui Rad, Bahram Masoudi

**Author notes:** **Corresponding authors:**, **1- Behnam Bakhshi**, Horticulture Crops Research Department, Sistan Agricultural and Natural Resources Research and Education Center, Agricultural Research, Education and Extension Organization (AREEO), Zabol, Iran, **2- Seyed Mohammad Taghi Tabatabaei**, Horticulture Crops Research Department, Yazd Agricultural and Natural Resources Research and Education Center, Agricultural Research, Education and Extension Organization (AREEO), Yazd, Iran.

## Abstract

Salinity is one of the most important causes of yield loss in agricultural products, especially wheat. Wheat cultivation, on the other hand, is carried out on a wide range of agricultural land in saline lands. Therefore, wheat breeding for tolerance to salinity can be an effective way to improve yield and yield stability under these conditions. In this study, twenty wheat genotypes were studied in a randomized complete block design with three replications in two normal and salinity stress conditions in order to identify suitable indices of wheat stress tolerance and also to identify genotypes tolerant to salinity stress. Genotypes were evaluated based on tolerance index (TOL), stress tolerance index (STI), mean productivity (MP), susceptibility to stress (SSI), geometric mean productivity (GMP), harmonic mean productivity (HM), yield stability (YS) and yield index (YS). The results showed that MP, GMP, HM and STI indices are suitable indices because of their positive and highly significant correlation with yield in both normal and salinity stress conditions and it was found that these indices were suitable tools to achieve high yield genotypes in both environments. The results also showed that genotypes 10, 4, 9, 3 and 8 are resistant to salt stress with acceptable yield. Genotypes 5, 11, 12, 14, 16, 17, 18 and 20 were also identified as the most susceptible genotypes.

## Introduction

Salinity is one of the main limiting factors in crop productivity, especially wheat. Salinity causes numerous problems due to induction of defective physiological functions of plant growth and development. Species and plant varieties are highly diverse in their ability to withstand salinity. Salt resistance varies even in species of the same genus (Francois et al. 1994). Wheat is a plant that due to its adaptation to different environmental conditions, cultivated in different regions of the world with diverse climatic conditions (Loss and Siddique 1994). The major production of wheat occurs in the latitudes of 30-60 degrees north and 27-40 degrees south, and in most of the major wheat producing countries, grain yield is important. Considering environmental conditions is one of the effective factors in selection and introduction of superior genotypes based on grain yield. Environmental stresses affect wheat growth from germination to late stages of growth (fertilization and grain filling). Stress during grain filling period decreases grain weight (Ehdaie and Waines 1996). Many traits such as spike length, number of grains per spike, and 1000-grain weight in wheat are affected by stress. The number of fertile spikelets and tillers is reduced, fertilization is disrupted and grain yield is eventually reduced (Maas and Grieve 1990; Fernandez 1992). Wheat growth decline is one of the plant responses to salinity stress that results from a decrease in the synthetic activities of cellular metabolic pathways that affect dry matter production and manifest as a decrease in plant leaf area and ultimately it affects the yield and growth of all plant organs (Maas and Grieve 1990). Breeding for salinity tolerance can be an effective way to improve yield and yield stability in arid soils (Genc et al. 2010). The purpose of producing salt tolerant and drought tolerant cultivars is to introduce cultivars that tolerate stress better than other genotypes and show less yield loss under identical conditions (Srivastava et al. 1987). In an experiment on wheat, it was found that salinity had no effect on spikelet production of wheat genotypes but shortened the duration of this step (Grieve et al. 1993). Dryness significantly reduced the number of leaves on the main stem and the number of spikelets per spike and greatly reduced the number of tillers (Passioura 2006). Increasing yield under water deficit conditions requires identification of resistant genotypes. Different quantitative indices have been proposed to evaluate the response of genotypes to environmental conditions and to determine their salinity and drought tolerance and sensitivity. In semi-arid regions with inadequate rainfall distribution, yield potential in stress condition is not the best criterion for stress resistance, but rather yield stability and comparison of yield under stress condition and normal conditions have been introduced for species response to moisture stress (Silim et al. 1988). The stress tolerance criterion is expressed as the grain yield in the stress condition (Fischer and Maurer 1978). Therefore, relative yield status of genotypes in stress condition and in appropriate condition is a starting point for identification of drought related traits and selection of genotypes for breeding in stressful environments (Golestani and Pakniyat 2007). Thus, the ability of the plant to produce in stress condition compared to yield in non-stress state has been proposed as a criterion of stress tolerance (Fischer and Maurer 1978). An appropriate selection index is an index that distinguishes high yield genotypes in both stress and non-stress conditions from other genotypes (Rosielle and Hamblin 1981). Rosielle and Hamblin (1981) introduced the tolerance index (TOL) and mean productivity (MP). High values of TOL index indicate genotype susceptibility to stress. Therefore, selection of genotypes is based on low TOL. Fischer and Maurer (1978) proposed the stress susceptibility index (SSI). Low SSI values indicate low yield variation of a genotype under stress and non-stress conditions. In other words, low SSI value indicates low yield variation of a genotype under stress and stability conditions. Fernandez (1992) used the stress tolerance index (STI) to identify genotypes that have high yield under both stress and non-stress conditions. The more stable genotypes based on this index have higher STI values. The high value of this index for the genotype indicates more drought resistance and more potential yield of the genotype. Another indicator introduced by Fernandez was the geometric mean productivity (GMP) (Fernandez 1992). Based on GMP index, yield of genotypes is calculated under stress and non-stress conditions. This index is less sensitive to different values of yield under normal and stress conditions (Fernandez 1992; Schneider et al. 1997). This index is more potent in separation of genotypes compared to MP. The lower the stress sensitivity index in the cultivar, the more tolerant it will be to stress conditions, and the closer the yield to normal and stress conditions, the greater the yield stability of the variety. Harmonic mean productivity (HM) was another indicator introduced by Schneider et al. (1997). Schneider et al. (1997) suggested that genotypes with higher HM values are more resistant. Another indicator is yield stability index (YSI) which indicates the amount of genetic resistance to stress (Bouslama and Schapaugh 1984). The genotype with high YSI index performs well in both stress and non-stress environments. Fernandez (1992) classified the plant into four groups by examining plant yield in two stress and non-stress environments: a) Genotypes that have the same expression in stress and non-stress environments (Group A), b) Genotypes that only exhibit good performance in a non-stress environment (Group B), c) High Yield Genotypes in stress environment (Group C), d) Genotypes with poor expression in both environments (Group D). According to Fernandez, the most appropriate criterion for selection is a criterion that can distinguish group A from other groups (Fernandez 1992). In the current study, twenty wheat genotypes will be evaluated for different salinity indices. Also, identifying the best tolerance indices and identifying salt tolerant genotypes were the other important goals of this study.

## Materials and Methods

In order to study stress tolerance indices and identification of salt tolerant genotypes in wheat, twenty wheat genotypes were studied in a randomized complete block design with three replications under two normal and salinity stress conditions at research station of Agricultural Research and Training Center and Natural resources of Yazd (Table 1). Water and soil EC were 2.5 and 3.5 dS / m in normal conditions and 11 and 12 dS / m in stress condition, respectively. Dimensions of each experimental plot were 3 m^2^ and each experimental plot in each block consisted of six 2.5 m planting lines with 20 cm^2^ distance. Irrigation of normal and saline plots varied in autumn and winter according to the region’s custom and according to the need of the plant, but in spring every 14 days, it was done according to the need of the plant in basin irrigation form. All planting and harvesting activities for both normal and stress conditions were the same and were done according to local customs and scientific methods. Then, quantitative indices of salinity tolerance for each genotype were calculated using seed yield of plants under stress and non-stress conditions. The formulas for the various indices are outlined in Table 2. Analysis of variance of quantitative indices of calculated indices for genotypes was done in a randomized complete block design. Correlation coefficients between yield and all indices were calculated under normal and stress conditions. Also, Fernandez graph and Fernandez drawing methods and biplot based on principal component analysis were used to identify superior genotypes and to investigate the relationship between genotypes and indices, thus, the most suitable and most resistant genotypes to salinity were determined. SPSS, Statgraphics and Excel were used in the current study for data analysis.

**Table 1:**
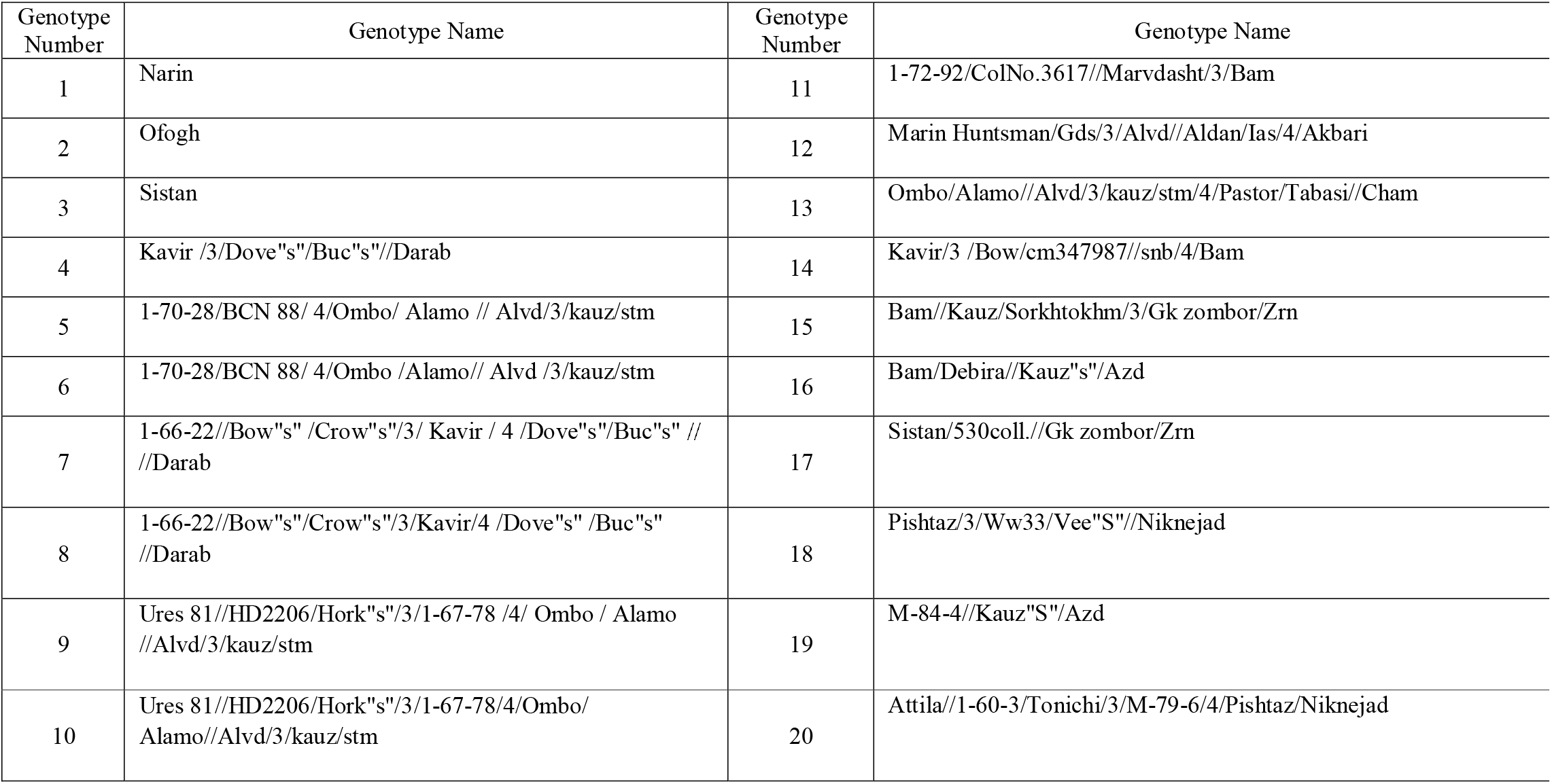
Evaluated genotype names and numbers

**Table 2:**
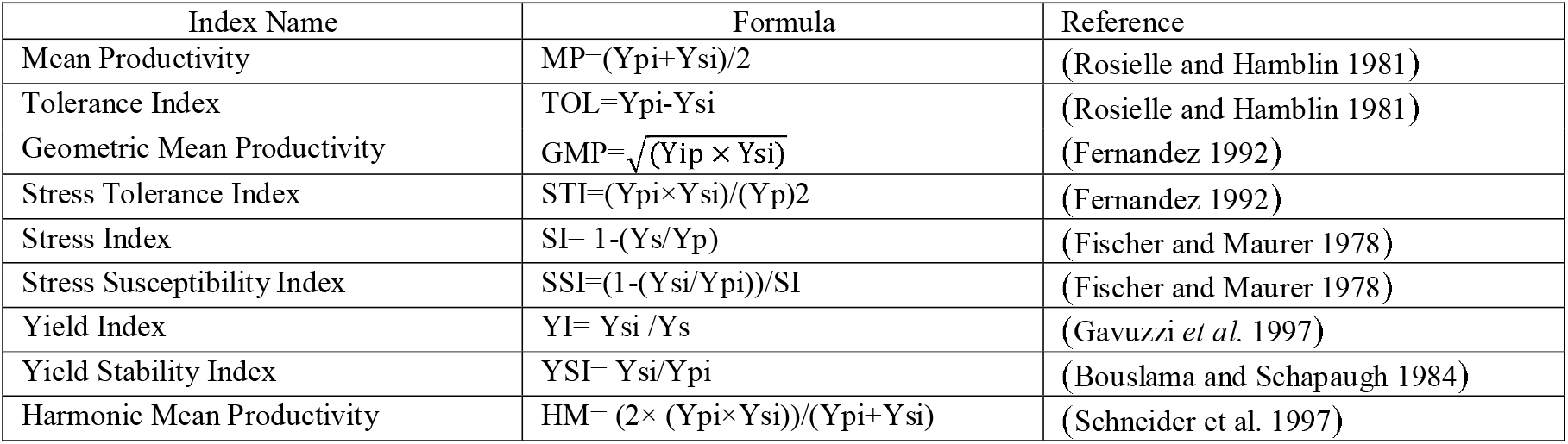
Indicators calculated in this study and their calculation method

| Index Name | Formula | Reference |
| --- | --- | --- |
| Mean Productivity | $MP = (Y_{pi} + Y_{si})/2$ | (Rosielle and Hamblin 1981) |
| Tolerance Index | $TOL = Y_{pi} - Y_{si}$ | (Rosielle and Hamblin 1981) |
| Geometric Mean Productivity | $GMP = \sqrt{(Y_{ip} \times Y_{si})}$ | (Fernandez 1992) |
| Stress Tolerance Index | $STI = (Y_{pi} \times Y_{si}) / (Y_p)^2$ | (Fernandez 1992) |
| Stress Index | $SI = 1 - (Y_s / Y_p)$ | (Fischer and Maurer 1978) |
| Stress Susceptibility Index | $SSI = (1 - (Y_{si} / Y_{pi})) / SI$ | (Fischer and Maurer 1978) |
| Yield Index | $YI = Y_{si} / Y_s$ | (Gavuzzi *et al.* 1997) |
| Yield Stability Index | $YSI = Y_{si} / Y_{pi}$ | (Bouslama and Schapaugh 1984) |
| Harmonic Mean Productivity | $HM = (2 \times (Y_{pi} \times Y_{si})) / (Y_{pi} + Y_{si})$ | (Schneider *et al.* 1997) |

## Results and discussion

The results of analysis of variance showed significant differences at 1% level between genotypes in terms of a large number of stress tolerance indices and grain yield under both normal and salinity stress conditions (Table 3) which indicates the existence of genetic variation and the possibility of selection for tolerance to salinity.

**Table 3:** Analysis of variance of quantitative indices of salt resistance in wheat genotypes

| Source of variation | Degree of freedom | Mean square |  |  |  |  |  |  |  |  |  |
| --- | --- | --- | --- | --- | --- | --- | --- | --- | --- | --- | --- |
|  |  | HM | YI | YSI | SSI | STI | GMP | MP | TOL | YiS | YiP |
| Block | 2 | 366384<sup>ns</sup> | 0.09* | 0.37<sup>ns</sup> | 308<sup>ns</sup> | 0.25<sup>ns</sup> | 317382<sup>ns</sup> | 270125<sup>ns</sup> | 2209796<sup>ns</sup> | 653907 * | 991241<sup>ns</sup> |
| Genotype | 19 | 644718** | 0.11** | 0.09<sup>ns</sup> | 73<sup>ns</sup> | 0.39** | 636202** | 630281** | 601242<sup>ns</sup> | 795244** | 765940** |
| Error | 38 | 160891 | 0.03 | 0.15 | 123 | 0.11 | 152816 | 148909 | 698519 | 188820 | 458258 |

In order to identify resistant genotypes, salinity indices were calculated for the studied genotypes (Table 4). According to STI, higher values of this index indicate more resistance of genotypes to stress. According to this index, genotypes 10, 4, 9, 3, 6 and 8 were more resistant to stress than other genotypes. The above mentioned genotypes had the highest values of STI index among the studied genotypes and were in high yield genotypes group in both normal and stress condition. Genotypes 18, 20, 12 and 17 were identified as the most susceptible genotypes based on this index. SSI index is more commonly used to remove susceptible genotypes and each genotype with higher values of this index is more susceptible to stress (Fischer and Maurer 1978). Based on this index and considering the mean of yield in both normal and stress conditions, genotypes 11, 15, 13, 3, 8, 12, 9, 14 and 4 were in good condition compared to other genotypes and had low recall index and were identified as the best genotypes. Genotypes 6, 1, 7, and 17 were identified as the most sensitive genotypes with the highest values. According to TOL resistance index, genotypes are more resistant to lower values of mentioned index (Rosielle and Hamblin 1981). According to this index, Genotypes 6, 1, 5, 7, 17, 18 and 20 with low values of this index were the highest salinity resistant genotypes compared to other genotypes. But genotypes 15 and 11 had the highest values of these indices and were therefore the most sensitive genotypes. According to the mean productivity (MP), the more resistant genotypes have higher values of this index (Rosielle and Hamblin 1981). Accordingly, genotypes 10, 4, 9, 3, 6 and 8 were the most resistant genotypes and genotypes 18, 20 and 11 with the least value of this index were the most susceptible genotypes. Based on GMP, genotypes 10, 9, 4, 3 and 8 were identified as the most resistant genotypes and genotypes 19, 20 and 11 were the most susceptible genotypes. Yield Stability Index (YSI) indicates the amount of genotype genetic resistance to salinity stress (Bouslama and Schapaugh 1984) and therefore high yield stability genotype should have high yield in both stress and non-stress environments. Based on this index, genotypes 25, 1, 2, 15 and 16 were the most resistant and genotypes 20, 18, 7, 17, 5, 1 and 6 were the most susceptible genotypes. Based on harmonic mean productivity, genotypes 6, 10, 2, 23, 31 and 19 were the most resistant and genotypes 28, 3, 9, 14 and 17 were identified as the most susceptible genotypes. Considering all the indices of susceptibility and resistance to salinity and yield in both normal and salinity conditions, genotypes 4, 9 and 10 were identified as superior and resistant to salinity and genotypes 18, 20 and 11 were susceptible to salt stress. Since genotype 10 is in good condition in stress and non-stress condition, it can be considered as the most suitable genotype for cultivation in stress and non-stress condition.

**Table 4:**
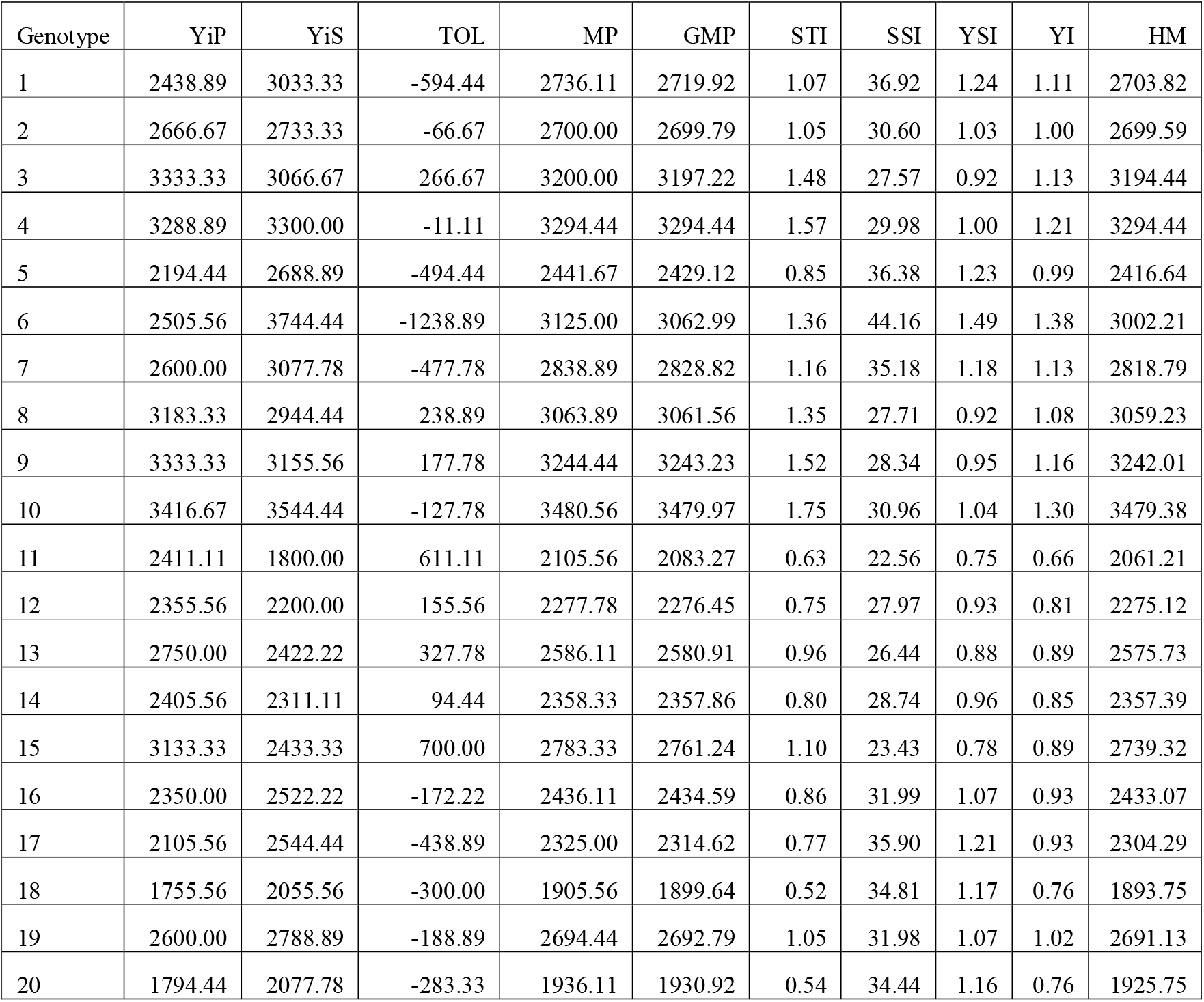
Estimation of drought tolerance indices based on average yield of wheat genotypes in stress and non-stress conditions

Simple correlation coefficients have been calculated and the results are shown in Fig. 1 to determine the appropriate indices and investigate the relationship between salt tolerant indices and yield under normal and stress conditions. According to Table 2, it was found that MP, GMP, HM and STI indices seemed to be suitable indices because of positive and highly significant correlation with yield in both normal and salinity conditions and selecting high values of these indices means high yield under normal conditions and salinity stress. All four indices showed positive and significant correlation with each other. According to Blum (2018), the best indicator should have a significant correlation with high yield in both normal and stress conditions. Therefore, given the significant and high correlation that MP, GMP, HM and STI indices had with yield in both conditions and with each other, they can be used to access high yield genotypes in both environments. There are several reports that introduced suitable indices for crops; including, HM, MP, STI and GMP indices for sesame genotypes; STI, MP and GMP indices for safflower genotypes; and STI and MP for bread wheat lines. These selections was based on significant differences between genotypes and species in terms of the mentioned indices which have enabled the study of genetic variation among genotypes (Usefiazar and Rezaei 2004; Golestani and Pakniyat 2007; Ganjeali et al. 2011). Also, in a study on chickpea, MP, GMP, STI and HM indices were selected as the best indices for selection of drought resistant genotypes (Ganjeali et al. 2011).

**Figure 1:**
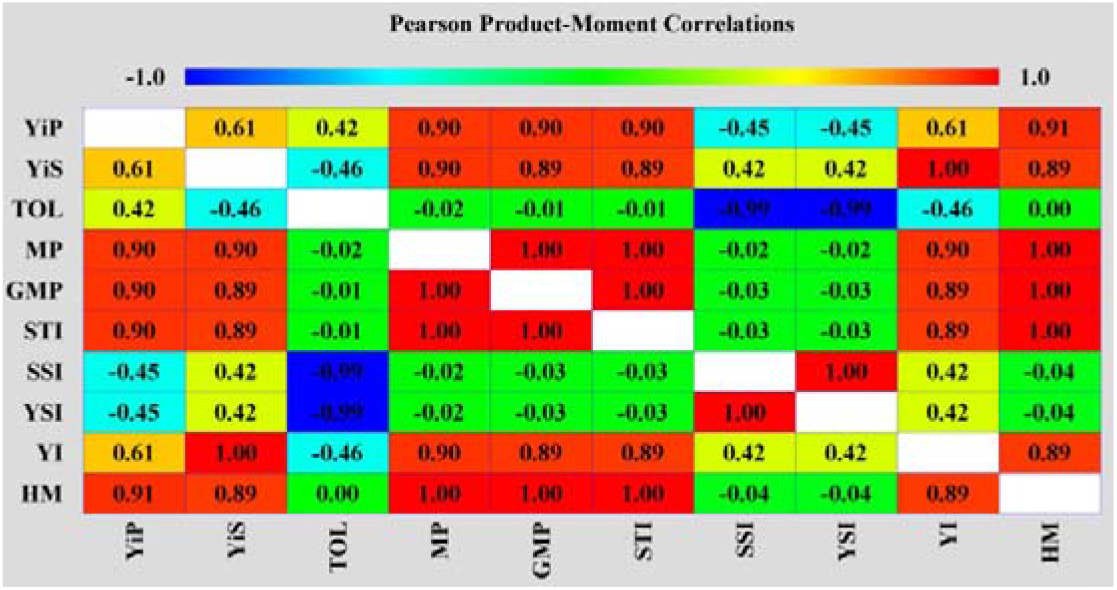
Correlation between salinity indices and grain yield

After identifying the best quantitative indices resistant to salt stress, three-dimensional diagrams were used to select high yielding genotypes in both stress and non-stress environments. This graph shows the relationship between the three variables Yip, Yis, and one of the resistance indices, where yield in the stress environment on the Y-axis, yield in the non-stress environment on the X-axis and the resistance indices on the Z-axis are shown. According to this criterion, genotypes are divided into four groups A, B, C and D; And according to Fernandez (1992), the most appropriate index is that the one that is able to distinguish group A from other groups. In this study, MP, GMP, HM and STI indices were identified as suitable indices for distinguishing group A from other groups, Therefore, their three-dimensional diagrams were used (Figures 2, 3, 4 and 5).

**Figure 2.**
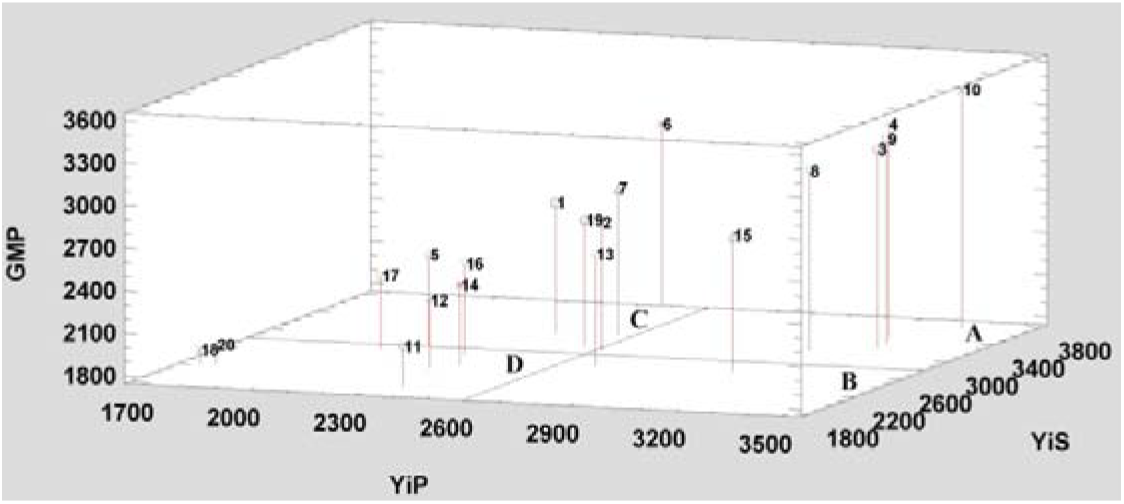
Three-Dimensional Distribution Diagram for Determination of Salinity Stress-resistant Wheat Genotypes Based on Yield Non-stress condition (YiP), yield stress condition (YiS) and GMP index

**Figure 3.**
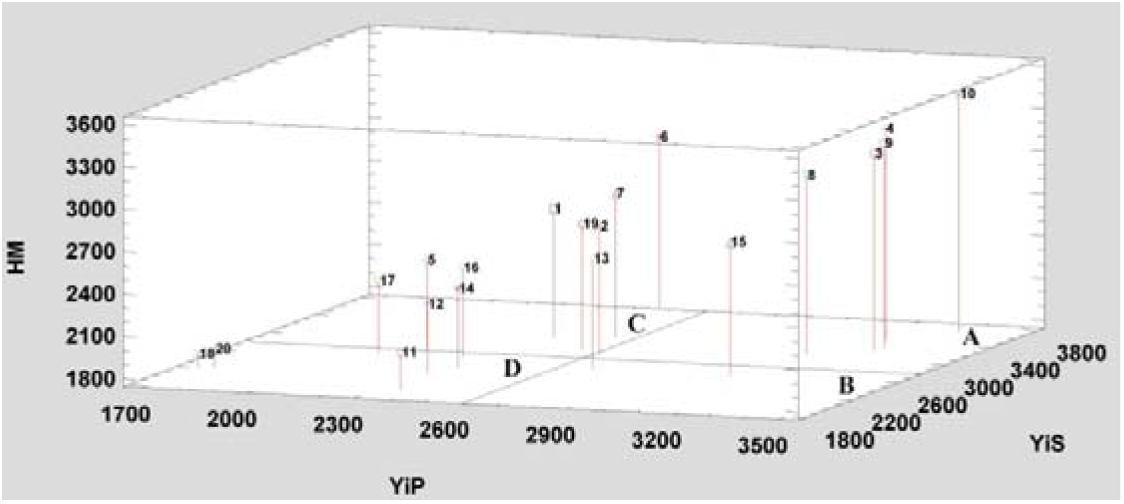
Three-Dimensional Distribution Diagram for Determination of Salinity Stress-resistant Wheat Genotypes Based on Yield Non-stress condition (YiP), yield stress condition (YiS) and HM Index

**Figure 4.**
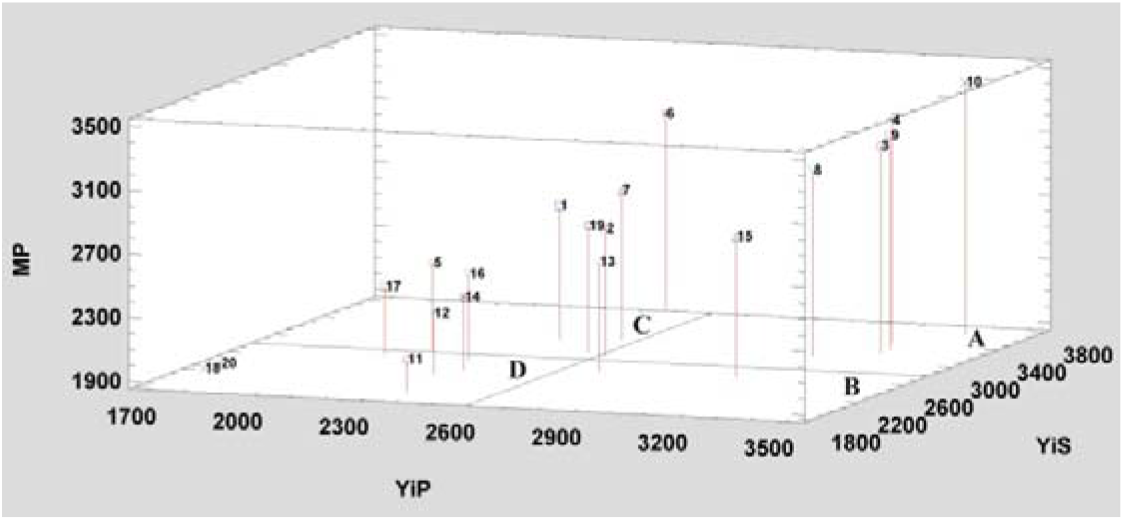
Three-Dimensional Distribution Diagram for Determination of Salinity Stress-resistant Wheat Genotypes Based on Yield Non-stress condition (YiP), yield stress condition (YiS) and MP Index

**Figure 5.**
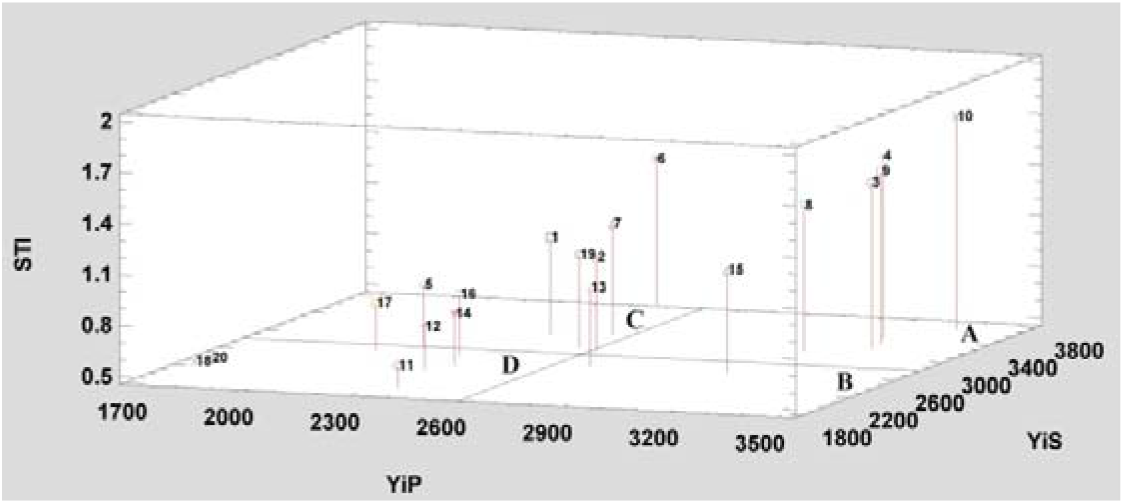
Three-Dimensional Distribution Diagram for Determination of Salinity Stress-resistant Wheat Genotypes Based on Yield Non-stress condition (YiP), yield stress condition (YiS) and STI Index

Based on the three-dimensional graph of Yis and Yip with MP, GMP, STI and HM indices, as can be seen, most of the genotypes were classified in A, C and D group. Genotypes 10, 4, 9, 3 and 8 belonged to group A and it means they are both resistant to salt stress and have high yields under normal conditions. Group C contains genotypes 1, 2, 5, 7, 16 and 19; Group D contains genotypes 11, 12, 14, 17, 18 and 20; and in group B, genotypes 13 and 15 were classified. The use of three-dimensional diagrams to identify Group A has been used and confirmed by Fernandez (1992) in beans and in chickpea by Ganjeali et al. (2011).

To investigate the relationship between more than three variables, a multivariate diagram called biplot is used. Thus, the relationships between the performance of genotypes and all resistance indices are shown in one figure. To plot this diagram, principal components analysis based on resistance and yield indices in normal and salinity stress conditions must first be performed. Factor coefficients were estimated based on principal component analysis method after Verimax rotation. According to Table 5, the most variation between the data with 99.8% was justified by the first two components. Therefore, the plot was drawn based on these two components. Also, this table shows that 64.52% of the total variation was related to the first component which had a positive correlation with YS, YP, MP, GMP, HM, YI and STI, and had a negative correlation with TOL and had a low correlation with SSI and YSI indices. Therefore, the first component was called potential and yield sustainability. Due to the high values of this component, selection of genotype with proper yield is possible under both normal and stress conditions. The second component, accounts for 35.28% of the total variance in the data, had negative correlation with yield in salinity stress condition and had negative and strong correlation with SSI and YSI and had high and positive correlation with TOL index. Therefore, the second component is highly correlated with salt sensitivity indices. For this reason, the second component was called the salt sensitivity component. Based on these two components, genotypes were divided in biplot space into specific groups based on mean yield and resistance to salt stress. Given the high correlation between yield and salinity stress with the first component and the positive correlation between the second component and salt sensitivity indices, genotypes that are located in the upper reaches of these two components are recommended as salt resistant with high yield. Therefore, based on the drawn biplot, genotypes 10, 4, 9, 3 and 8 located in high salinity potential and low susceptibility to salinity and adjacent to vectors related to important salinity indices such as MP, GMP HM and STI, were identified as high yielding resistant genotypes. Genotypes 1, 2, 7, 13, 15 and 19, which were in low yield area under high stress and susceptibility to salinity and adjacent to important salt sensitivity indices TOL and SSI, were identified as genotypes with specific adaptations to different environments. Additionally, genotypes 5, 11, 12, 14, 16, 17, 18 and 20, located in low yield area under normal and salinity stress conditions were identified as low-yielding genotypes in both conditions. Since the angle between the vectors shows the degree of correlation between the variables, the sharp angle between the MP, GMP, HARM and STI indices indicates a strong correlation between these indices. The results of this diagram (Fig. 6) confirm the results of the three-dimensional diagrams of Figures 1, 2, 3 and 4. The use of principal component analysis and bioplot diagrams to differentiate genotypes for drought stress in beans has been used and confirmed by Fernandez (1992) and in chickpea by Ganjeali et al. (2011).

**Table 5:** Eigenvalues, Eigenvectors and cumulative contribution of resistance and yield indices of wheat genotypes in stress and non-stress conditions

| Component | Cumulative portion Percent | Eigenvalues | HM | YI | YSI | SSI | STI | GMP | MP | TOL | YiS | YiP |
| --- | --- | --- | --- | --- | --- | --- | --- | --- | --- | --- | --- | --- |
| 1 | 64.52 | 6.45 | 0.39 | 0.37 | 0.04 | 0.04 | 0.39 | 0.39 | 0.39 | -0.05 | 0.37 | 0.33 |
| 2 | 99.80 | 3.53 | 0.07 | -0.18 | -0.53 | -0.53 | 0.07 | 0.07 | 0.06 | 0.53 | -0.18 | 0.29 |

**Figure 6.**
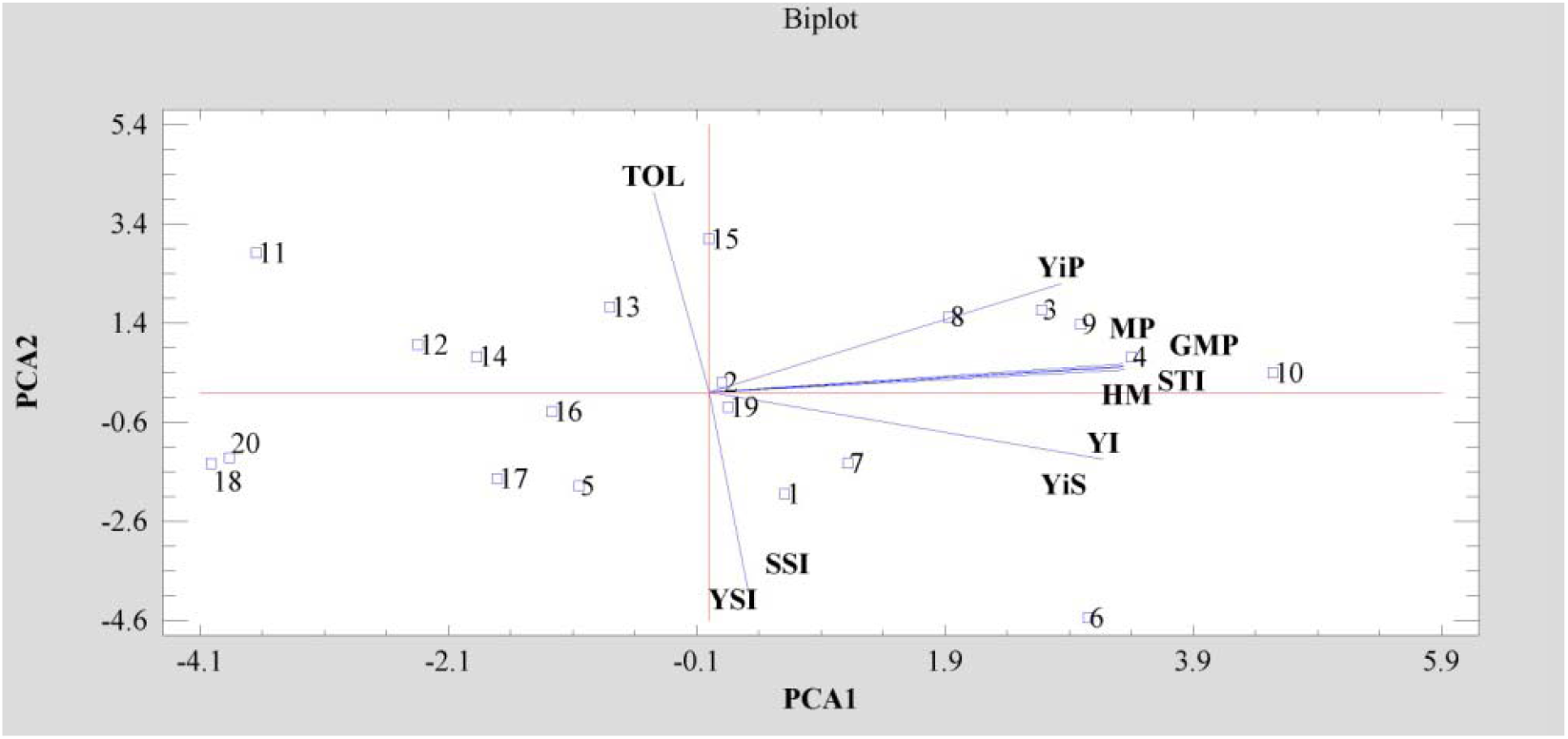
Biplot figure of wheat genotypes for 8 tolerant indices based on two first components.

## Conclusion

Different wheat genotypes investigated in this experiment showed high variation in grain yield as well as resistance, susceptibility and response to salinity indices. Based on the results of correlation analysis of resistance indices with grain yield in stress and non-stress condition, STI, MP, GMP and HM indices were the best indices in wheat for selection of salt tolerant genotypes. Finally, according to the set of studies, salinity resistant wheat genotypes 10, 4, 9, 3 and 8 are recommended.

---

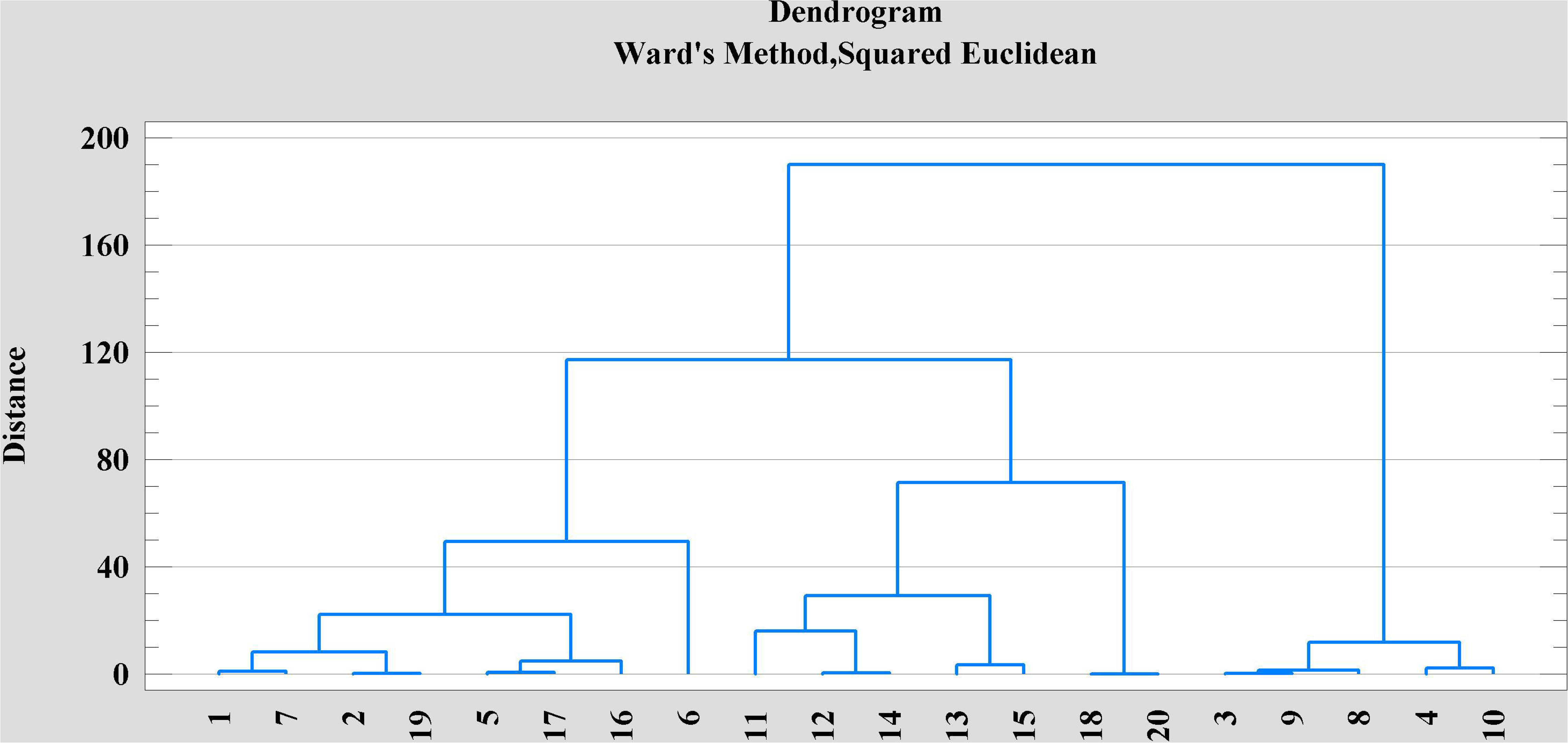

## Notes

### Competing Interest Statement

The authors have declared no competing interest.

